# Itaconate and fumarate derivatives exert a dual inhibitory effect on canonical NLRP3 activation in macrophages and microglia

**DOI:** 10.1101/2021.02.01.429180

**Authors:** Christopher Hoyle, Jack P Green, Stuart M Allan, David Brough, Eloise Lemarchand

## Abstract

The NLRP3 inflammasome is a multi-protein complex that regulates the protease caspase-1 and subsequent interleukin (IL)-1β release from cells of the innate immune system, or microglia in the brain, in response to infection or injury. Derivatives of the metabolites itaconate and fumarate, dimethyl itaconate (DMI), 4-octyl itaconate (4OI) and dimethyl fumarate (DMF), limit both expression of IL-1β, and IL-1β release following NLRP3 inflammasome activation. However, the direct effects of these metabolite derivatives on NLRP3 inflammasome responses in macrophages and microglia require further investigation. Using murine bone marrow-derived macrophages, mixed glia and organotypic hippocampal slice cultures (OHSCs), we demonstrate that DMI and 4OI pre-treatment limited IL-1β, IL-6 and tumor necrosis factor production in response to lipopolysaccharide (LPS) priming, as well as inhibiting subsequent NLRP3 inflammasome activation. DMI, 4OI, DMF and monomethyl fumarate (MMF), another fumarate derivative, also directly inhibited biochemical markers of NLRP3 activation in LPS-primed macrophages, mixed glia and OHSCs, including ASC speck formation, caspase-1 activation, gasdermin D cleavage and IL-1β release. Finally, DMF, an approved treatment for multiple sclerosis, as well as DMI, 4OI and MMF, inhibited NLRP3 activation in macrophages in response to the phospholipid lysophosphatidylcholine, which is used to induce demyelination, suggesting a possible mechanism of action for DMF in multiple sclerosis through NLRP3 inhibition. Together, these findings reveal the importance of immunometabolic regulation for both the priming and activation steps of NLRP3 activation in macrophages and microglia. Furthermore, we highlight itaconate and fumarate derivatives as a potential therapeutic option in NLRP3-driven diseases, including in the brain.

**Summary statement:** We show that itaconate and fumarate derivatives inhibit both the priming and activation steps of NLRP3 inflammasome responses in macrophages and microglia, revealing the importance of immunometabolic NLRP3 regulation.

## 1 Introduction

Macrophages are innate immune effector cells that regulate inflammatory responses upon infection or tissue injury to restore tissue homeostasis by promoting pathogen death or tissue and wound repair. In the brain, macrophage-like cells called microglia are important effectors of this inflammatory response. Inflammasomes are cytosolic complexes that regulate the inflammatory response in immune cells and microglia. In particular, the nucleotide-binding oligomerisation domain-, leucine-rich repeat- and pyrin domain-containing protein 3 (NLRP3) inflammasome has been implicated in a range of non-communicable diseases that are characterised by an inflammatory response (Chen and Nuñez, 2010; Rock et al., 2010). Although several pathways of NLRP3 activation have been described (Gaidt et al., 2016; Kayagaki et al., 2011), canonical NLRP3 activation is the most studied. The canonical pathway consists of an initial priming step, typically mediated through Toll-like receptor signaling, that upregulates NLRP3 and IL-1β expression, followed by a subsequent NLRP3 activating stimulus. A broad range of pathogen- or damage-associated molecular patterns are known to act as this activating stimulus, including the potassium ionophore nigericin, extracellular ATP (Perregaux and Gabel, 1994), amyloid-β aggregates (Halle et al., 2008) or silica crystals (Dostert et al., 2008; Hornung et al., 2008). The precise mechanism by which these stimuli induce NLRP3 activation is still unclear, with potassium efflux-dependent (Muñoz-Planillo et al., 2013) and -independent (Groß et al., 2016) mechanisms suggested to elicit dispersal of the trans-Golgi network, leading to inflammasome formation (Chen and Chen, 2018). We recently proposed organelle dysfunction to be a crucial cellular event that leads to NLRP3 activation (Seoane et al., 2020). Once activated, NLRP3 interacts with the adaptor protein ASC (apoptosis-associated speck-like protein containing a caspase recruitment domain), causing the formation of an ASC speck that drives activation of the inflammasome effector protein caspase-1 (Boucher et al., 2018; Schroder and Tschopp, 2010). Active caspase-1 then cleaves gasdermin D and pro-interleukin (IL)-1β, with gasdermin D pores potentially forming the conduit for mature IL-1β release (He et al., 2015).

Immunometabolism has emerged as a regulator of macrophage inflammasome responses (O’Neill and Artyomov, 2019). Lipopolysaccharide (LPS) treatment of macrophages causes a metabolic shift from oxidative phosphorylation to glycolysis that is necessary for IL-1β production (Tannahill et al., 2013). Certain metabolites such as itaconate (O’Neill and Artyomov, 2019), succinate (Mills and O’Neill, 2014) and fumarate (Humphries et al., 2020) have immunoregulatory functions. For example, itaconate and fumarate derivatives, including dimethyl itaconate (DMI), 4-octyl itaconate (4OI) and dimethyl fumarate (DMF), are able to activate nuclear factor erythroid 2-related factor 2 (NRF2) signalling by alkylating and subsequently inducing the degradation of the cytoplasmic NRF2 inhibitor kelch-like ECH-associated protein 1 (KEAP1) (Mills et al., 2018). NRF2 is then able to translocate to the nucleus, where it not only upregulates the transcription of its target genes, but also prevents the recruitment of RNA polymerase II to NF-κB secondary response genes such as IL-6 and IL-1β (Kobayashi et al., 2016). DMI and 4OI also induce electrophilic stress and glutathione depletion in macrophages, which inhibits the LPS-induced translation of IκBζ independently of NRF2, and this subsequently limits the expression of IκBζ-dependent NF-κB secondary response genes (Bambouskova et al., 2018; Swain et al., 2020). It must be acknowledged that the properties of these itaconate derivatives may not fully reflect the properties of endogenous itaconate (Swain et al., 2020). *In vivo* evidence also indicates the importance of itaconate responses, as mice deficient in *Irg1*, which therefore cannot produce itaconate, rapidly succumb to *Mycobacterium tuberculosis* infection, whereas there was no mortality in wild-type control mice (Nair et al., 2018). Interestingly, 4OI and DMF exhibit anti-viral and anti-inflammatory effects through NRF2 signalling in response to Severe Acute Respiratory Syndrome Coronavirus 2 (SARS-CoV2) infection (Olagnier et al., 2020).

Although itaconate derivatives are known to limit IL-1β expression, the direct effects of itaconate-related compounds on NLRP3 inflammasome activation are less characterized. Previous studies suggest that DMF, an approved treatment for relapsing-remitting multiple sclerosis, and its metabolite monomethyl fumarate (MMF), limit NLRP3 inflammasome activation, with DMF exhibiting greater potency (Liu et al., 2016; Miglio et al., 2015). DMF has also been shown to directly succinate a cysteine residue on gasdermin D to limit pyroptotic cell death in response to NLRP3 activation *in vitro* and *in vivo*, but without inhibiting NLRP3 activation itself (Humphries et al., 2020). Itaconate and its derivative 4OI can inhibit NLRP3 activation in an NRF2-independent manner, through the modification of specific cysteine residues on NLRP3, which may prevent NLRP3’s interaction with NEK7 and subsequent activation (Hooftman et al., 2020; Swain et al., 2020). IRG1-deficient macrophages, which cannot synthesize endogenous itaconate, exhibit enhanced IL-1β release in response to NLRP3 inflammasome activation. Finally, 4OI is effective at inhibiting NLRP3 activation *in vivo* (Hooftman et al., 2020).

Despite recent advances, further characterisation of the effects of itaconate- and fumarate-related compounds on NLRP3 inflammasome activation is required in order to evaluate their therapeutic potential. The relevance of immunometabolic regulation of microglial inflammasome responses in the brain is also unclear. Here, we demonstrate that itaconate derivative pre-treatment not only prevented expression of IL-1β, but also inhibited canonical NLRP3 inflammasome activation. We identified that itaconate and fumarate derivatives were able to directly inhibit canonical NLRP3 inflammasome activation, independent of their inhibitory effect on priming. These effects were consistent in mixed glia and organotypic hippocampal slice cultures (OHSCs), two brain-relevant NLRP3 inflammasome models (Hoyle et al., 2020). Finally, itaconate and fumarate derivatives inhibited NLRP3 activation induced by lysophosphatidylcholine (LPC), a lipid molecule used to induce demyelination in models of multiple sclerosis, further highlighting a potential mechanism of DMF action in multiple sclerosis treatment. These findings reveal a dual anti-inflammatory effect of itaconate and fumarate derivatives in the innate immune system and brain, through regulation of both the priming and activation steps of canonical NLRP3 inflammasome responses.

## 2 Results

### Pre-treatment with itaconate and fumarate derivatives reduces NLRP3 priming and canonical inflammasome activation

BMDMs were pre-treated with two cell-permeable derivatives of itaconate, DMI and 4OI, as well as the fumarate derivative DMF, and the effects on NLRP3 priming and activation were assessed. DMI and 4OI treatment alone did not induce NRF2 accumulation in murine WT BMDMs, but both enhanced LPS-induced NRF2 accumulation (Figure 1A). DMF treatment was toxic to the cells at this treatment duration and concentration, explaining the lack of NRF2 accumulation (Figure 1A, Supplementary Figure 1). IκBζ protein levels were not measured. DMI and 4OI pre-treatment inhibited the production of pro-IL-1β in response to LPS priming, and whilst DMI reduced NLRP3 expression to a lesser extent, 4OI had no effect on NLRP3 levels (Figure 1A). No pro-IL-1β or NLRP3 protein was observed following DMF pre-treatment, although this was probably due to toxicity prior to LPS priming (Figure 1A, Supplementary Figure 1). DMI and 4OI also strongly reduced LPS-induced IL-6 release, with a smaller reduction in TNF release (Figure 1Bi, Bii). Consistent with inhibition of pro-IL-1β expression, DMI and 4OI pre-treatment blocked IL-1β release in response to subsequent stimulation with LPS and nigericin (Figure 1Biii). These data confirmed previous findings that itaconate derivative treatment prior to LPS stimulation could limit inflammatory priming via NRF2 activation (Mills et al., 2018).

**Figure 1.**
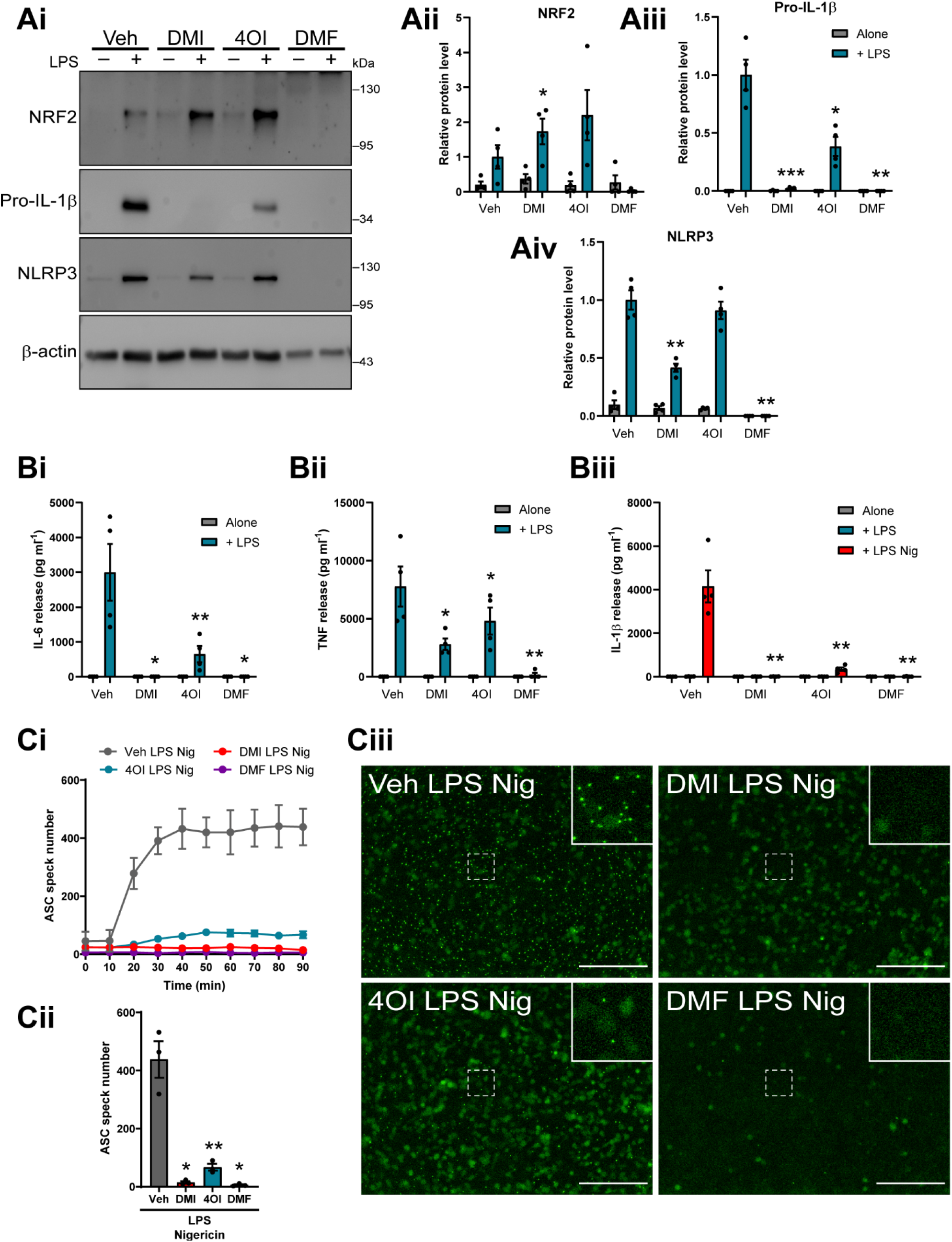
Pre-treatment with itaconate and fumarate derivatives reduces NLRP3 priming and canonical inflammasome activation. (**A**) WT BMDMs were treated with vehicle (DMSO), DMI, 4OI or DMF (125 μM, 20 h). LPS (1 μg ml^−1^, 4 h) was then added to the wells to induce priming (n=4). (**Ai**) Cell lysates were probed by western blotting for NRF2, pro-IL-1β and NLRP3 protein, and (**Aii– iv**) densitometry was performed on each independent experiment (expressed relative to Veh+LPS treatment). (**B**) WT BMDMs were treated as above, followed by nigericin (10 μM, 60 min; n=4). Supernatants were assessed for (**Bi**) IL-6, (**Bii**) TNF and (**Biii**) IL-1β content by ELISA. (**C**) ASC– citrine BMDMs were treated as above, followed by nigericin (10 μM, 90 min; n=3). ASC speck formation was measured over a period of 90 min. Image acquisition began immediately after addition of nigericin. (**Ci**) ASC speck number per field of view was quantified over 90 min. (**Cii**) ASC speck number and (**Ciii**) fluorescence images after 90 min nigericin treatment are shown. Scale bars are 200 μm. Data are presented as mean ± SEM. Data were analysed using repeated-measures one-way (Cii) or two-way (Aii–iv, B) ANOVA with Dunnett’s post-hoc test (versus Veh treatment within each group). *P<0.05; **P<0.01, ***P<0.001.

We next investigated whether itaconate derivative pre-treatment could reduce NLRP3 inflammasome activation, as has been recently suggested (Swain et al., 2020). Treatment of murine BMDMs from ASC–citrine reporter mice (Tzeng et al., 2016) with DMI, 4OI and DMF prior to LPS priming inhibited the formation of ASC specks upon subsequent nigericin treatment (Figure 1Ci, Cii), with representative images from this experiment after 90 minutes of nigericin treatment shown (Figure 1Ciii). Morphological changes due to nigericin treatment are also shown using bright-field microscopy (Supplementary Figure 1). DMF pre-treatment induced a high level of cell toxicity, perhaps explaining its strong inhibitory effect on NLRP3 inflammasome formation and IL-1β release. These data suggested that itaconate derivative pre-treatment may additionally inhibit the NLRP3 inflammasome activation step, as well as inhibiting the priming stage.

### Itaconate and fumarate derivatives directly inhibit the NLRP3 activation step

To determine whether NLRP3 inflammasome inhibition was a direct effect of the itaconate derivatives, LPS-primed WT BMDMs were treated with DMI, 4OI and DMF prior to nigericin stimulation, and this resulted in inhibition of IL-1β release, as well as small reductions in cell death (Figure 2Ai, Aii). Western blotting of the cell lysates demonstrated that the levels of pro-IL-1β were consistent between treatments, confirming that in this protocol expression of pro-IL-1β was unaffected by DMI, 4OI or DMF (Figure 2Aiii, Supplementary Figure 2A). Dose-dependent inhibition of IL-1β release was observed for each treatment, with minimal reductions in cell death (Supplementary Figure 3). ASC speck formation was measured in LPS-primed ASC–citrine BMDMs treated with DMI, 4OI and DMF prior to nigericin. Each of these metabolite derivatives inhibited ASC speck formation in response to nigericin, suggesting that these compounds were also able to directly block NLRP3 inflammasome activation independently of their effects on the priming response (Figure 2Bi–iii). Fluorescence and bright-field images are shown (Figure 2Bi, Supplementary Figure 4). DMF treatment after LPS priming did not induce morphologically observable cell death, indicating that DMF was able to inhibit NLRP3 inflammasome activation independent of toxicity at this concentration and duration of incubation, and this has been reported previously (Garstkiewicz et al., 2017; Liu et al., 2016; Miglio et al., 2015) (Supplementary Figure 3). LPS-primed primary BMDMs treated with DMI, 4OI or DMF and subsequent nigericin stimulation were lysed directly in-well without removing the supernatant, and western blotting confirmed reductions in caspase-1 activation, and gasdermin D and IL-1β cleavage (Figure 2C, Supplementary Figure 2B). NRF2 levels were increased by DMI, 4OI and DMF treatment after LPS priming, although this was not observed in cells that received subsequent nigericin stimulation (Figure 2C, Supplementary Figure 2B), and itaconate-mediated NLRP3 inhibition is suggested to be independent of NRF2 (Hooftman et al., 2020; Swain et al., 2020). MMF treatment limited NLRP3 activation in LPS-primed BMDMs, although it was not as potent as DMF (Fig 2Di, Dii). We confirmed that exogenous itaconate treatment also inhibited NLRP3 activation in LPS-primed BMDMs, although much higher doses were required because it is less cell permeable (Supplementary Figure 5) (Swain et al., 2020). To determine whether the inhibitory effects of itaconate and fumarate derivatives were relevant in human macrophages, LPS-primed human MDMs were treated with DMI, 4OI and DMF prior to nigericin stimulation. 4OI and DMF alone appeared to induce IL-1β release, although slight increases in cell death were observed for these treatments, which may have allowed passive release of unprocessed pro-IL-1β (Figure 2Ei). Both DMI and DMF reduced nigericin-induced IL-1β release, whereas 4OI did not significantly reduce IL-1β release at this dose (Figure 2Ei). No inhibition of cell death was observed (Figure 2Eii). Thus, the itaconate and fumarate derivatives were able to directly inhibit NLRP3 activation in peripheral macrophages, independent of their effects on priming.

**Figure 2.**
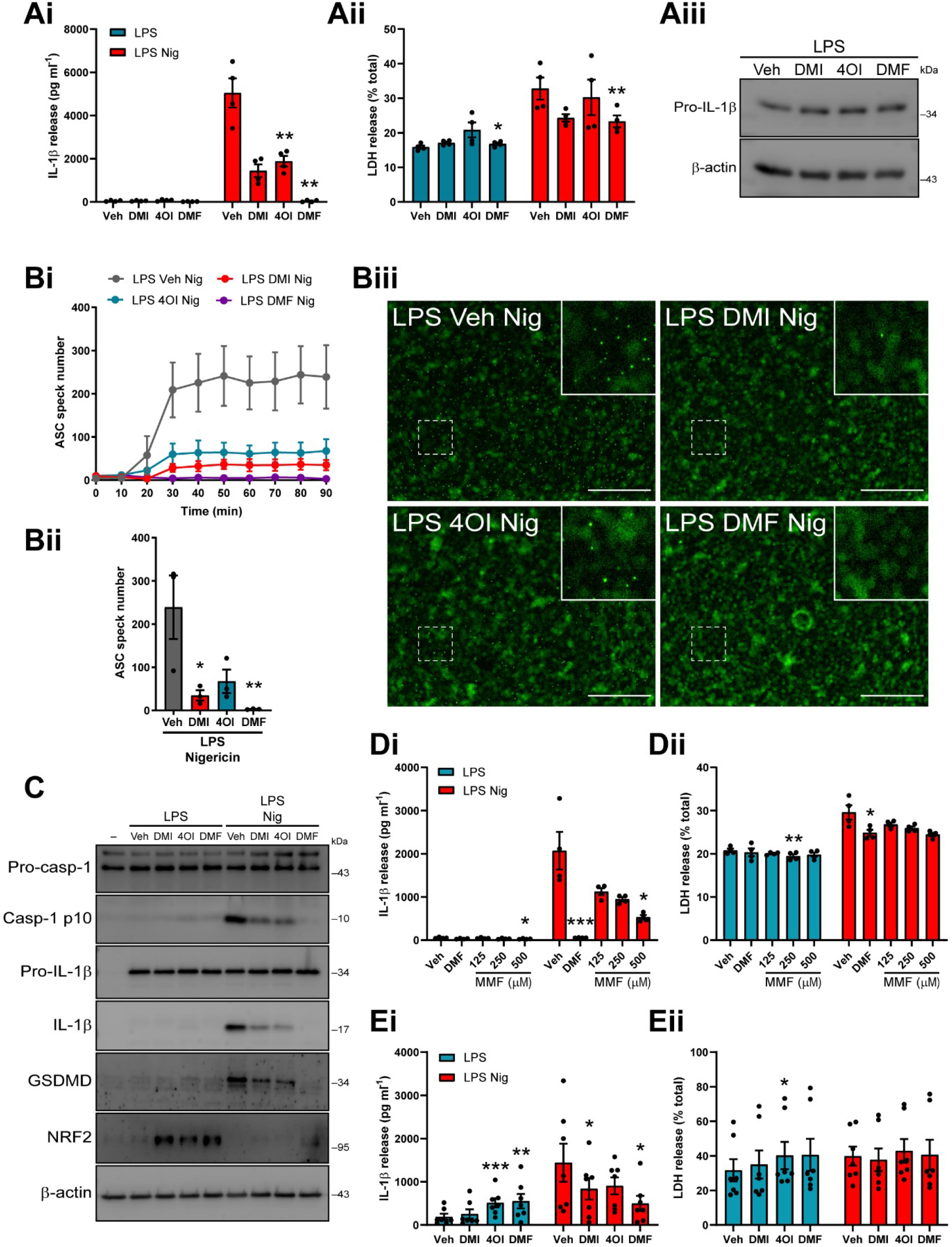
Itaconate and fumarate derivatives directly inhibit the NLRP3 activation step in murine and human macrophages. (**A**) WT BMDMs were primed with LPS (1 μg ml^−1^, 4 h) before treatment with vehicle (DMSO), DMI, 4OI or DMF (125 μM, 15 min). Nigericin was then added to the well (10 μM, 60 min; n=4). Supernatants were assessed for (**Ai**) IL-1β release and (**Aii**) cell death (LDH release). (**Aiii**) Cell lysates were probed by western blotting for pro-IL-1β protein. See Supplementary Figure 2A. (**B**) ASC–citrine BMDMs were treated as above, and ASC speck formation was measured over a period of 90 min (n=3). Image acquisition began immediately after addition of nigericin. (**Bi**) ASC speck number per field of view was quantified over 90 min. (**Bii**) ASC speck number and (**Biii**) fluorescence images after 90 min nigericin treatment are shown. Scale bars are 200 μm. (**C**) WT BMDMs were treated as above and then lysed in-well and probed for several markers of inflammasome activation by western blotting (n=4). See Supplementary Figure 2B. (**D**) WT BMDMs were LPS primed (1 μg ml^−1^, 4 h) before treatment with vehicle (DMSO), DMF (125 μM) or MMF (125–500 μM, 15 min). Nigericin was then added to the well (10 μM, 60 min; n=4). Supernatants were assessed for (**Di**) IL-1β release and (**Dii**) LDH release. **(E**) Human MDMs were LPS primed (1 μg ml^−1^, 4 h) before treatment with vehicle (DMSO), DMI, 4OI or DMF (125 μM, 15 min). Nigericin was then added to the well (10 μM, 60 min; n=7). Supernatants were assessed for (**Ei**) IL-1β release and (**Eii**) LDH release. Supernatants were assessed for cytokine content by ELISA. Data are presented as mean ± SEM. Data were analysed using repeated-measures one-way (Bii) or two-way (Ai, Aii, D, E) ANOVA with Dunnett’s post-hoc test (versus Veh treatment within each group). *P<0.05; **P<0.01; ***P<0.001.

### Itaconate and fumarate derivatives inhibit NLRP3 priming and canonical activation in mixed glia and OHSCs

Despite accumulating evidence in peripheral immune cells, little is known about the importance of immunometabolic regulation of inflammasome priming and activation in the brain, where microglia are thought to be the predominant source of inflammasomes (Sheppard et al., 2019). Thus, we assessed the effect of itaconate derivative pre-treatment on mixed glial cultures, which consist of approximately 80% astrocytes, 10% microglia and 10% oligodendrocyte/type-2 astrocyte progenitor cells (Pinteaux et al., 2002), and organotypic hippocampal slice cultures (OHSCs), which we recently validated as a model for studying microglial NLRP3 responses (Hoyle et al., 2020). DMF was not included in these pre-treatment experiments due to its toxicity observed in murine BMDMs (Figure 1, Supplementary Figure 1), although it has been previously demonstrated to inhibit IL-1β, IL-6, and to a lesser extent TNF expression at a lower dose in rat neonatal microglial cultures (Wilms et al., 2010). DMI and 4OI alone did not induce detectable NRF2 accumulation in mixed glial cultures (Figure 3Ai, Aii). DMI treatment did not significantly enhance LPS-induced NRF2 accumulation, although increases were observed; similarly, 4OI did not increase LPS-induced NRF2 levels (Figure 3Ai, Aii). IκBζ protein levels were not measured. Despite this, both DMI and 4OI reduced pro-IL-1β production in mixed glial cultures, without affecting NLRP3 protein levels (Figure 3Ai–iv). IL-1β release upon subsequent stimulation with nigericin was also inhibited by both derivatives, and 4OI modestly reduced cell death (Figure 3Av, Avi). Given that the inhibition of pro-IL-1β production and mature IL-1β release was comparable between DMI and 4OI in mixed glia, only 4OI was used in OHSCs. NRF2 accumulation could not be reliably detected in OHSCs upon LPS priming (data not shown). 4OI reduced the production of pro-IL-1β in response to LPS priming, but did not affect NLRP3 production (Figure 3Bi– iii). 4OI pre-treatment strongly inhibited IL-1β release in response to LPS and nigericin treatment and reduced cell death (Figure 3Biv, Bv). These data suggested that itaconate derivatives were able to limit the priming of inflammasome responses, and that this may be a relevant mechanism to regulate microglial inflammatory gene expression.

**Figure 3.**
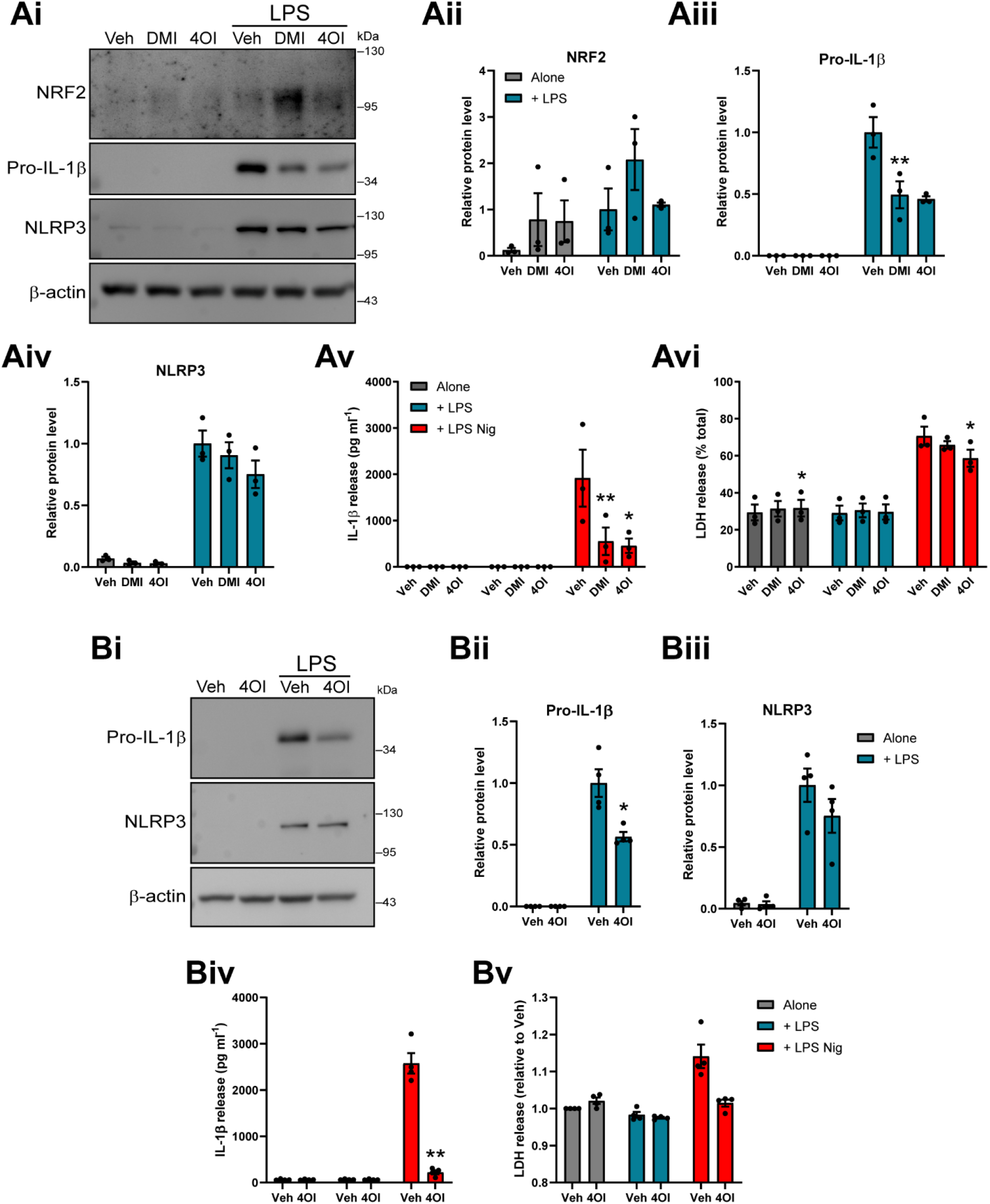
Itaconate and fumarate derivative pre-treatment inhibits NLRP3 priming in mixed glia and OHSCs. (**A**) WT mixed glia were treated with vehicle (DMSO), DMI or 4OI (125 μM, 21 h). LPS (1 μg ml^−1^, 3 h) was then added to the wells to induce priming (n=3). (**Ai**) Cell lysates were probed by western blotting for NRF2, pro-IL-1β and NLRP3 protein, and (**Aii–iv**) densitometry was performed on each independent experiment (expressed relative to Veh + LPS treatment). (**Av, Avi**) WT mixed glia were treated as above, followed by nigericin (10 μM, 60 min; n=3). Supernatants were assessed for (**Av**) IL-1β release and (**Avi**) cell death (LDH release). (**B**) WT OHSCs were treated with vehicle (DMSO) or 4OI (125 μM, 21 h). LPS (1 μg ml^−1^, 3 h) was then added to the wells to induce priming (n=4). (**Bi**) OHSC lysates were probed by western blotting for pro-IL-1β and NLRP3 protein, and (**Bii– iiii**) densitometry was performed on each independent experiment (expressed relative to Veh+LPS treatment). (B**iv, Bv**) WT OHSCs were treated as above, followed by nigericin (10 μM, 90 min; n=4). Supernatants were assessed for (**Biv**) IL-1β release and (**Bv**) LDH release. Supernatants were assessed for cytokine content by ELISA. Data are presented as mean ± SEM. Data were analysed using repeated-measures two-way ANOVA with Dunnett’s (A) or Sidak’s (B) post-hoc test (versus Veh treatment within each group). *P<0.05; **P<0.01; ***P<0.001.

Given that 4OI appeared to inhibit mature IL-1β release more strongly than it inhibited pro-IL-1β production in OHSCs, we investigated whether itaconate and fumarate derivatives could directly limit NLRP3 inflammasome activation in mixed glia and OHSCs. LPS-primed mixed glial cultures were treated with DMI and 4OI prior to nigericin stimulation, and this reduced IL-1β release but did not inhibit cell death (Figure 4Ai, Aii). Similarly, LPS-primed OHSCs were treated with DMI, 4OI, DMF and MMF prior to nigericin stimulation. The compounds alone did not exhibit any toxicity, although a small increase in cell death was observed in response to DMF treatment, nor did they induce ASC speck formation (Supplementary Figure 6). Following nigericin stimulation, 4OI and DMF significantly inhibited IL-1β release, though this was not accompanied by significant reductions in ASC speck formation (Figure 4Bi, Bii). Only MMF treatment significantly reduced nigericin-induced cell death, although this reduction was modest (Figure 4Biii). Representative immunofluorescence images are shown (Figure 4Biv). Together, these data suggested that the itaconate and fumarate derivatives could reduce microglial NLRP3 responses.

**Figure 4.**
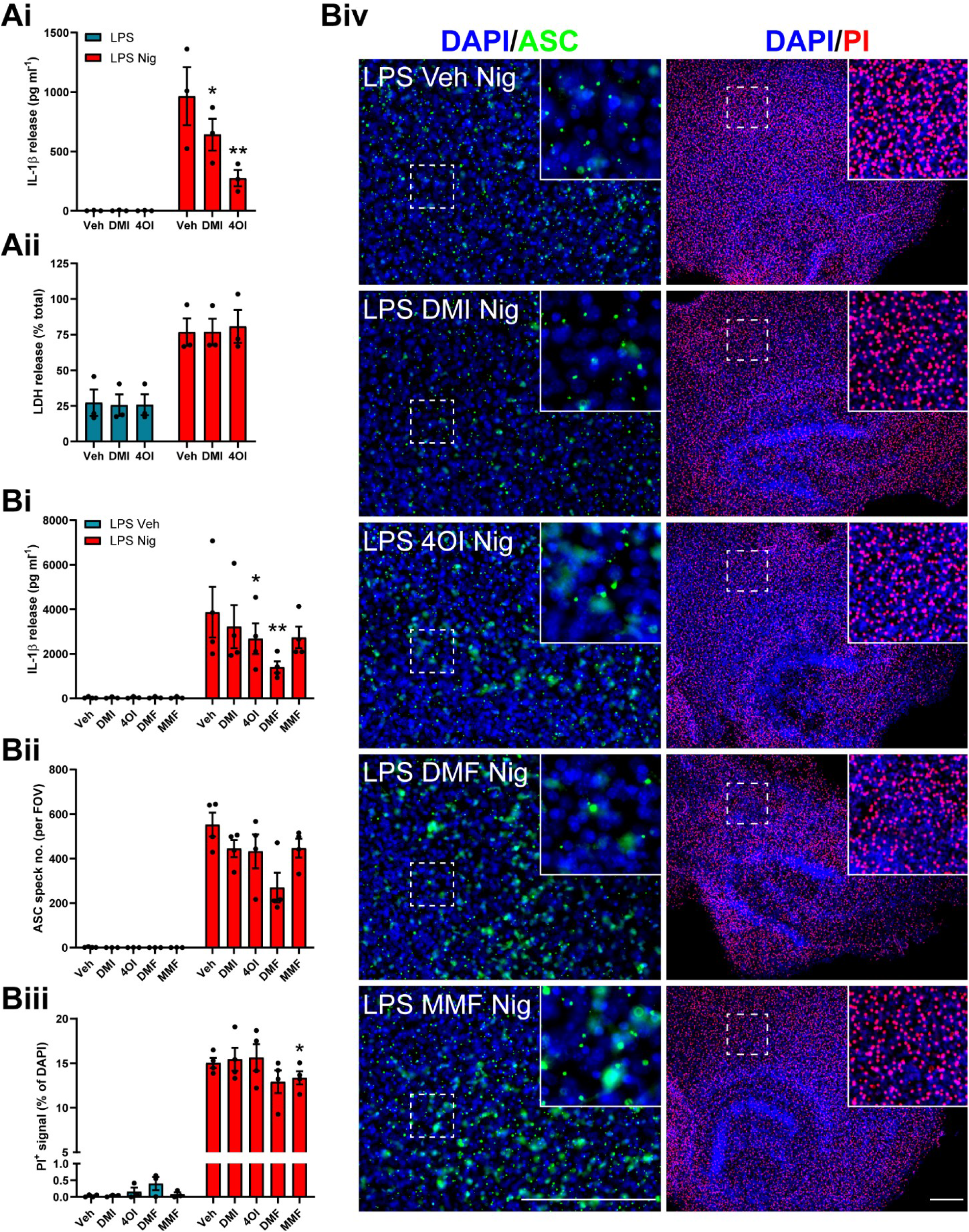
Itaconate and fumarate derivatives partly inhibit canonical NLRP3 activation in LPS-primed mixed glia and OHSCs. (**A**) Mixed glia were primed with LPS (1 μg ml^−1^, 3 h) before treatment with vehicle, DMI or 4OI (125 μM, 15 min). Nigericin was then added to the well (10 μM, 60 min; n=3). Supernatants were assessed for (**Ai**) IL-1β release and (**Aii**) cell death (LDH release). (**B**) WT OHSCs were primed with LPS (1 μg ml^−1^, 3 h) before treatment with vehicle (DMSO), DMI, 4OI, DMF (125 μM) or MMF (500 μM, 15 min). Vehicle (ethanol) or nigericin was then added to the well (10 μM, 90 min; n=3–4). Propodium iodide (PI; red, 25 μg ml^−1^) was added for the final 30 min of nigericin treatment. (**Bi**) Supernatants were assessed for IL-1β content. OHSCs were probed for nuclei (DAPI, blue) and ASC (green) by immunofluorescence staining. (**Bii**) ASC speck number per field of view was quantified. (**Biii**) The area of PI-positive staining was determined and is expressed as a % of total area of DAPI staining. (**Biv**) Representative images are shown. Images were acquired using widefield microscopy at 20X (ASC) and 5X (PI) magnification. Scale bars are 200 μm. Data are presented as mean ± SEM. Data were analysed using repeated-measures two-way ANOVA (A) or mixed effects model (B) with Dunnett’s post-hoc test (versus Veh treatment within each group). *P<0.05; **P<0.01.

### Itaconate and fumarate derivatives inhibit NLRP3 activation in response to LPC stimulation

LPC (also known as lysolecithin) is a lipid biomolecule that is generated from the cleavage of phosphatidylcholine by phospholipase A2, or via the action of lecithin-cholesterol acyltransferase (Law et al., 2019). LPC levels are regulated by the enzyme lysophosphatidylcholine acyltransferase, which converts LPC back to phosphatidylcholine (Law et al., 2019). Despite its presence during normal physiology, LPC is able to induce demyelination in experimental models of multiple sclerosis (Hall, 1972; Lassmann and Bradl, 2017; Plemel et al., 2018), implicating endogenous LPC dysregulation as a potential factor in multiple sclerosis pathology, although evidence of phospholipase A2 involvement in multiple sclerosis patients is unclear (Trotter et al., 2019). LPC can also activate the NLRP3 and NLRC4 inflammasomes in macrophages, microglia and astrocytes (Freeman et al., 2017), and NLRP3 is reported to be detrimental in experimental autoimmune encephalomyelitis (Coll et al., 2015; Gris et al., 2010; Jha et al., 2010). Thus, LPC-induced NLRP3 inflammasome activation may directly influence LPC-induced demyelination. We assessed whether the itaconate and fumarate derivatives could inhibit NLRP3 activation driven by LPC stimulation in peripheral macrophages. LPS-primed BMDMs were treated with DMI, 4OI, DMF or MCC950, a selective NLRP3 inhibitor (Coll et al., 2015), prior to LPC stimulation. Each of the derivatives inhibited IL-1β release to the same extent as MCC950, indicating inhibition of NLRP3 activation, although no reductions in cell death were observed by any treatment (Figure 5Ai, Aii). Similarly, treatment with MMF inhibited IL-1β release induced by LPC to the same extent as MCC950 (Figure 5Bi, Bii). These data suggest that itaconate and fumarate derivatives could limit NLRP3 activation in peripheral macrophages in response to LPC stimulation.

**Figure 5.**
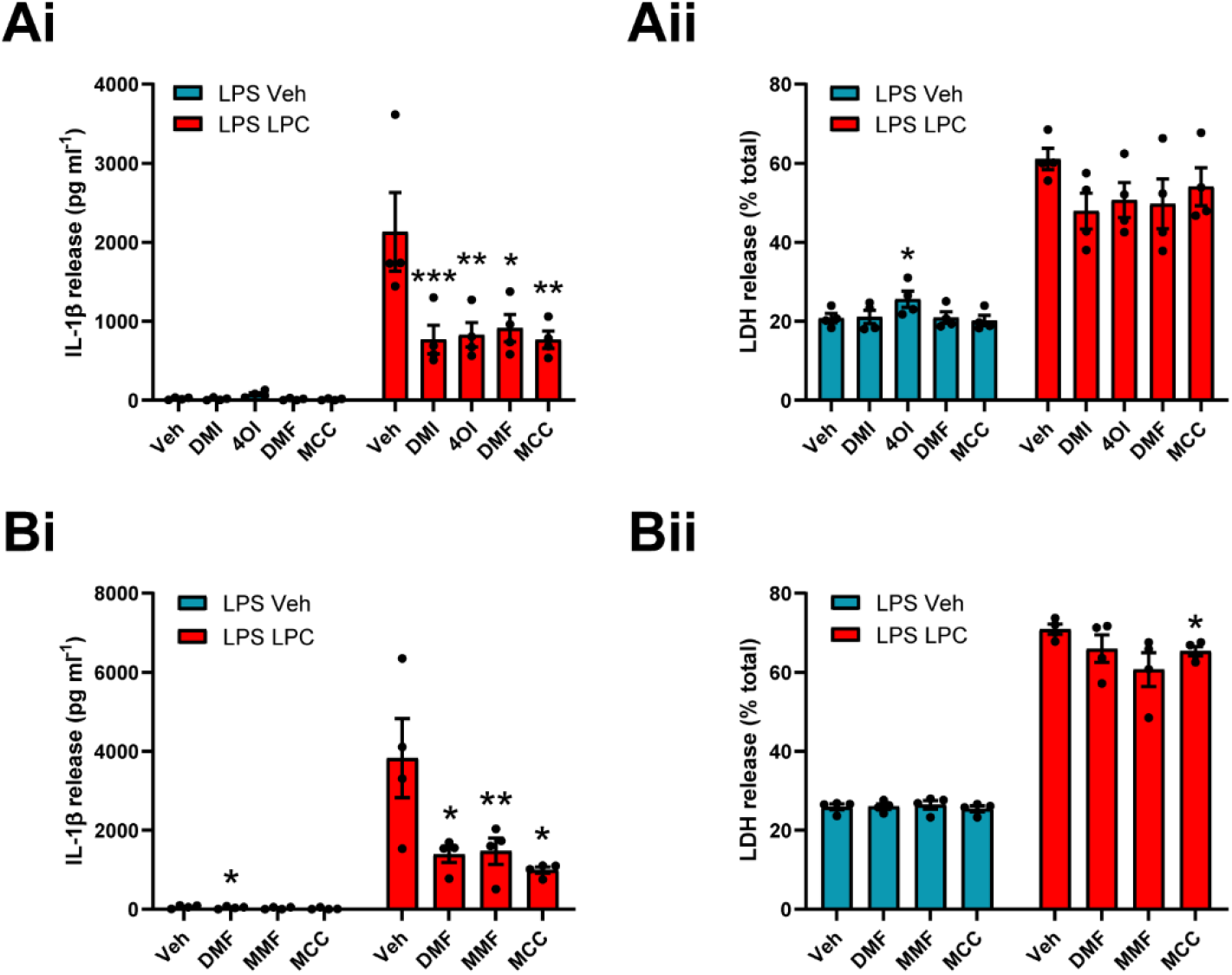
Itaconate and fumarate derivatives inhibit NLRP3 activation in response to LPC stimulation. (**A**) WT BMDMs were primed with LPS (1 μg ml^−1^, 4 h) before treatment with vehicle (DMSO), DMI, 4OI, DMF (125 μM) or MCC950 (10 μM, 15 min). Vehicle (ethanol) or LPC (100 μM, 60 min) was then added to the well (n=4). (**B**) WT BMDMs were LPS primed (1 μg ml^−1^, 4 h) before treatment with vehicle (DMSO), DMF (125 μM), MMF (500 μM) or MCC950 (10 μM, 15 min). Vehicle (ethanol) or LPC (100 μM, 60 min) was then added to the well (n=4). Supernatants were assessed for (**Ai, Bi**) IL-1β release by ELISA and (**Aii, Bii**) cell death (LDH release). Data are presented as mean ± SEM. Data were analysed using repeated-measures two-way ANOVA with Dunnett’s post-hoc test (versus Veh treatment within each group). *P<0.05; **P<0.01; ***P<0.001.

## 3 Discussion

Macrophage metabolism has emerged as an important regulator of inflammatory responses (O’Neill and Artyomov, 2019). In particular, exogenous treatment with derivatives of the metabolite itaconate inhibits the expression of NF-κB secondary response genes such as IL-1β and IL-6 through NRF2 accumulation (Mills et al., 2018) and inhibition of IκBζ translation (Bambouskova et al., 2018). However, the direct effect of itaconate derivative treatment on NLRP3 inflammasome activity is unclear, with a few recent studies indicating inhibition of IL-1β release independently of reductions in pro-IL-1β (Hooftman et al., 2020; Swain et al., 2020). We confirmed that DMI and 4OI are able to inhibit the expression of pro-inflammatory genes in macrophages upon LPS priming and show that this mechanism may have relevance in microglia. We also found that DMI, 4OI, DMF and MMF can directly inhibit biochemical hallmarks of NLRP3 inflammasome activation independently of their effects on inflammasome priming, including in human macrophages. These findings highlight that itaconate and fumarate derivatives could potentially be manipulated therapeutically in NLRP3-driven diseases, conferring the dual benefit of targeting both priming and activation of NLRP3.

Exogenous itaconate has different properties from commonly used itaconate derivatives such as DMI and 4OI, which exhibit greater electrophilicity (Swain et al., 2020). For example, exogenous itaconate does not appear to strongly drive NRF2 signalling, nor does it inhibit IκBζ translation, and so any effects of itaconate derivatives, although still relevant, may not directly reflect the physiology of endogenous itaconate (Swain et al., 2020). It has also been shown that DMI is not metabolised to itaconate in the cytosol (ElAzzouny et al., 2017). Nevertheless, whilst itaconate derivatives may not be appropriate to simulate the actions of endogenous itaconate, any therapeutic potential for these compounds maintains importance. Interestingly, Swain et al. (2020) also suggested that itaconate derivatives, including exogenous itaconate, can inhibit NLRP3 activation independently of effects on NRF2 and priming. However, itaconate treatments were only applied prior to LPS priming, instead of treating LPS-primed BMDMs with itaconate prior to NLRP3 activation. Thus, it remains possible that other effects of itaconate pre-treatment, such as reduction in NLRP3 protein levels during LPS priming, could have limited IL-1β release. We have addressed this in the current study, complementing observations that 4OI specifically inhibited NLRP3 activation in LPS-primed cells through direct interaction with cysteine 548 on murine NLRP3, preventing NEK7 binding (Hooftman et al., 2020). This mechanism is plausible, given that DMI, 4OI and DMF are electrophiles that modify cysteine residues on target proteins including KEAP1 (Linker et al., 2011; Mills et al., 2018), GAPDH (Kornberg et al., 2018; Liao et al., 2019), and more than a thousand other proteins in LPS-primed RAW macrophages (Qin et al., 2020).

We suggest that the inhibitory mechanisms of itaconate and fumarate derivatives on NLRP3 priming and canonical activation may be consistent in microglia. Thus, the implications of immunometabolic regulation of inflammasome responses in the brain should be explored further, particularly in the context of brain pathology. Transcriptomic databases indicate that microglia exhibit relatively high expression of *Nfe2l2* (NRF2) and *Keap1* (Schaum et al., 2018; Zhang et al., 2014). The itaconate-synthesising enzyme *Irg1* (also known as *Acod1*) was recently shown to be upregulated in response to LPS treatment in OHSCs (Chausse et al., 2020), driving subsequent itaconate production, and this was prevented by microglial depletion, indicating that this is primarily a microglial response. Furthermore, exogenous treatment with 4OI limited LPS-induced IL-6 production but did not affect TNF (Chausse et al., 2020). While mixed glia and OHSCs are useful tools to study microglial function, further studies are required to fully understand the physiological relevance of microglial metabolic reprogramming *in vivo*. NRF2 activation is associated with a protective effect in experimental ischaemic stroke models (Liu et al., 2019), and IRG-deficient mice exhibit exacerbated brain damage to acute ischaemic stroke, according to a recent pre-print (Kuo et al., 2020b). These studies suggest that itaconate production could be an endogenous, protective response to limit ischaemic damage. Our previous report showed increased levels of IL-1β and NLRP3 expression after ischaemic stroke, but NLRP3 deficiency or inhibition did not improve stroke outcome (Lemarchand et al., 2019). This could suggest that endogenous itaconate production within the brain in response to ischaemia, whilst too late to inhibit inflammatory cytokine production, may be able to limit NLRP3 inflammasome activation. It is important to note that detection of increased NRF2 levels in response to DMI and 4OI treatment was not reliable in the microglial models employed in this study, either due to a lower amount of NRF2 signalling, questioning the relevance of NRF2 activation in the brain, or due to the instability of NRF2 protein during OHSC sample processing steps such as water-bath sonication. It should also be noted that the extent of NLRP3 inhibition mediated by the itaconate and fumarate derivatives in the OHSCs was lower than in the BMDM and mixed glial assays. It is possible that higher doses or longer treatment times of the metabolite derivatives would result in greater NLRP3 inhibition, given that we have previously shown that MCC950 is able to potently inhibit IL-1β release and ASC speck formation in OHSCs (Hoyle et al., 2020).

We demonstrate that DMF, an approved clinical treatment for relapsing-remitting multiple sclerosis (Fox et al., 2012; Gold et al., 2012b; Linker et al., 2011) and psoriasis (Smith, 2017), is an effective NLRP3 inhibitor in both macrophages and microglia, as has been suggested previously (Garstkiewicz et al., 2017; Liu et al., 2016; Miglio et al., 2015). We also show that DMF, as well as DMI and 4OI, limits NLRP3 activation in response to LPC stimulation of macrophages, a demyelinating agent that can drive NLRP3 and NLRC4 activation in macrophages, microglia and astrocytes (Freeman et al., 2017). The mechanism underlying DMF’s beneficial effect in multiple sclerosis and psoriasis is unclear, and is commonly suggested to be mediated via NRF2 activation, although NRF2-independent effects are also reported (Schulze-Topphoff et al., 2016). Given that DMF is able to potently inhibit NLRP3 activation and directly inhibit gasdermin D cleavage (Humphries et al., 2020), and that NLRP3 is detrimental in the experimental autoimmune encephalomyelitis model of multiple sclerosis (Coll et al., 2015; Gris et al., 2010; Jha et al., 2010), it is possible that DMF’s protective response in multiple sclerosis is in part mediated through dampened microglial and macrophage NLRP3 activation that may promote or result from neuronal demyelination. Inhibition of NLRP3 activation in macrophages or microglia could also facilitate other inflammatory responses that promote clearance of damaged myelin and remyelination (Cunha et al., 2020). Evidence of caspase-1 activation and gasdermin D-mediated pyroptosis has also been observed in the CNS of multiple sclerosis patients and in animal models (Li et al., 2019; McKenzie et al., 2018), further implicating NLRP3 involvement. Upon administration, DMF is hydrolysed to MMF and can be detected in the serum (Gold et al., 2012a; Litjens et al., 2004). MMF also induces KEAP1 cysteine alkylation and NRF2 activation (Linker et al., 2011), suggesting it may exert similar inhibitory effects to DMF on the priming response. Importantly, here we confirm that MMF can inhibit NLRP3 activation in response to both nigericin and LPC stimulation in macrophages, suggesting that NLRP3 inhibition could indeed be a relevant *in vivo* mechanism for DMF treatment. While DMF exhibited toxic effects during the pre-treatment experiments, as its dose was matched to that of 4OI, it is likely that titration of the DMF concentration would result in reduced toxicity but similar potency. Given that DMI and 4OI, which both also drive NRF2 accumulation, exerted similar inhibitory effects on nigericin- and LPC-induced NLRP3 activation, it is possible that these itaconate derivatives may also offer therapeutic potential in the treatment of multiple sclerosis. Indeed, DMI was recently shown to be protective in a mouse model of multiple sclerosis (Kuo et al., 2020a).

We have revealed a two-pronged immunometabolic mechanism of NLRP3 regulation by limiting both NLRP3 priming and canonical activation, suggesting that treatment with derivatives of metabolites such as itaconate and fumarate may represent a viable therapeutic strategy in NLRP3-driven diseases, although further work is required to confirm this. Future work should also aim to establish whether itaconate and fumarate derivatives inhibit NLRP3 activation through the same or differing mechanisms, either confirming this as a promising therapeutic target for drug design, or revealing novel targets.

## 4 Materials and methods

### Mice

In-house colonies of wild-type (WT) and ASC–citrine (Tzeng et al., 2016) C57BL/6 mice at the University of Manchester were maintained to provide primary cell cultures. Animals were allowed free access to food and water and maintained under temperature-, humidity- and light-controlled conditions. All animal procedures adhered to the UK Animals (Scientific Procedures) Act (1986).

### Primary murine BMDM preparation

Primary bone marrow-derived macrophages (BMDMs) were prepared by centrifuging the femurs of 3–6-month-old WT or ASC–citrine mice of either sex in an Eppendorf tube containing phosphate-buffered saline (PBS) at 10,000 × g (10 s). Bone marrow was collected and red blood cells were lysed with ACK lysing buffer (Lonza, LZ10-548E). Cells were passed through a cell strainer (70 μm pore size; Corning, 734-2761), centrifuged at 1500 × g (5 min), and BMDMs were generated by resuspending and culturing the cell pellet in 70% Dulbecco’s modified Eagle’s medium (DMEM; Sigma, D6429) containing 10% (v/v) foetal bovine serum (FBS; Thermo, 10500064), 100 U ml^−1^ penicillin and 100 μg ml^−1^ streptomycin (PenStrep; Thermo, 15070063), and supplemented with 30% L929 mouse fibroblast-conditioned medium for 7 days. Cells were incubated at 37°C, 90% humidity and 5% CO2. Before experiments, BMDMs were seeded overnight at a density of 1 × 10^6^ ml^−1^.

### Human monocyte-derived macrophage preparation

Human monocyte-derived macrophages (MDMs) were prepared from human peripheral blood mononuclear cells (PBMCs) obtained from consenting healthy donors (National Health Service Blood and Transplant, Manchester, UK), with full ethical approval from the University Research Ethics Committee at the University of Manchester (ref 2017-2551-3945). In brief, PBMCs were isolated by Ficoll separation (Thermo) at 400 × g (40 min, room temperature) with zero deceleration. PBMCs were washed three times with sterile MACS buffer (0.5% (w/v) bovine serum albumin (BSA), 2 mM EDTA in PBS) before positive selection of CD14^+^ monocytes by incubation with magnetic CD14 microbeads (Miltenyi Biotec, 130-050-201) (15 min, 4°C) and elution using LS columns (Miltenyi Biotec, 130-042-401). CD14^+^ monocytes were differentiated to MDMs by culturing for 7 days (at a concentration of 1 × 10^6^ cells ml^−1^) in RPMI 1640 (Sigma, R8758) supplemented with 10% (v/v) FBS, PenStrep and macrophage colony-stimulating factor (M-CSF, 0.5 ng ml^−1^; Peprotech, 300-25) at 37°C, 90% humidity and 5% CO2. On day 3 of differentiation, cells were fed by the addition of fresh media containing M-CSF (0.5 ng ml^−1^). Before experiments, MDMs were seeded overnight at a density of 1 × 10^6^ ml^−1^.

### Primary murine mixed glial culture preparation

Murine mixed glial cells were prepared from the brains of 2–4-day-old mice of either sex that were killed by cervical dislocation, as described previously (Hoyle et al., 2020). The brains were isolated, cerebral hemispheres dissected and the meninges removed. The remaining brain tissue was homogenised in DMEM containing 10% (v/v) FBS and PenStrep via repeated trituration, then centrifuged at 500 × g for 10 min and the pellet was resuspended in fresh culture medium before being incubated in a flask at 37°C, 90% humidity and 5% CO_2_. After 5 days, the cells were washed, and fresh medium was applied. The medium was subsequently replaced every 2 days. On day 12 of the culture, the cells were seeded at 2 × 10^5^ cells ml^−1^ in 24-well plates and incubated for a further 2 days prior to use.

### Organotypic hippocampal slice culture (OHSC) preparation

Seven-day-old mouse pups of either sex were killed by cervical dislocation and the brains were collected in PBS containing glucose (5 mg ml^−1^). The hippocampi were dissected and placed on filter paper, and 400 μm slices were prepared using a McIlwain tissue chopper (Brinkman Instruments). Hippocampal slices were collected and placed on 0.4 μm Millicell culture inserts (Merck Millipore, PICM03050), as described previously by Stoppini et al. (1991). Three hippocampal slices were placed on each insert. Slices were maintained in a humidified incubator with 5% CO_2_ at 37°C with 1 ml MEM (Gibco, 31095209) containing 20% (v/v) horse serum (Sigma, H1138), supplemented with HEPES (30 mM; Fisher, 10397023) and insulin (0.1 mg ml^−1^; Gibco, 12585014), pH 7.2–7.3. The culture medium was changed every 2 days and slices were used at day 7.

### Treatment protocols

To assess the effect of itaconate and fumarate derivative pre-treatments on inflammasome priming, cells were first treated with vehicle (DMSO), DMI (Sigma, 592498), 4OI (Cayman Chemical, CAY25374) or DMF (all 125 μM; Sigma, 242926) for 20 h (BMDM) or 21 h (mixed glia and OHSC). LPS (1 μg ml^−1^; Sigma, L2654) was then added to the wells for 4 h (BMDM) or 3 h (mixed glia and OHSC) to induce priming, followed by nigericin (10 μM; Sigma, N7143) for 60 min (BMDM and mixed glia) or 90 min (OHSC) to activate the NLRP3 inflammasome.

To assess the direct effect of itaconate and fumarate derivative treatments on canonical NLRP3 inflammasome activation in macrophages, BMDMs and human MDMs were first primed with LPS (1 μg ml^−1^) for 4 h. The medium was then replaced with serum-free DMEM (BMDM) or RPMI (human MDM) containing vehicle (DMSO), DMI, 4OI, DMF (all 125 μM, 15 min), MMF (500 μM, 15 min Sigma, 651419), exogenous itaconate (1–7.5 mM, 15 min; Sigma, I29204) or the NLRP3 inhibitor MCC950 (10 μM, 15 min; Sigma, PZ0280), before nigericin (10 μM, 60 min) or LPC (100 μM in ethanol, 60 min; Sigma, L4129) was added to the culture medium. To assess the direct effect of itaconate and fumarate derivative treatments on NLRP3 inflammasome activation in mixed glia and OHSCs, cells were first primed with LPS (1 μg ml^−1^) for 3 h. The medium was then replaced with serum-free DMEM (mixed glia) or MEM (OHSC) containing vehicle (DMSO), DMI, 4OI, DMF (125 μM, 15 min) or MMF (500 μM, 15 min), before nigericin (10 μM) was added to the culture medium for 60 min (mixed glia) or 90 min (OHSC). At the end of the experiments, the supernatants were collected and cell or OHSC lysates prepared for further analysis.

### Western blotting

Primary BMDMs, mixed glia and OHSCs were lysed with lysis buffer (50 mM Tris/HCl, 150 mM NaCl, Triton-X-100 1% v/v, pH 7.3) containing protease inhibitor cocktail (Merck Millipore, 539131). OHSCs were additionally lysed using repeated trituration and brief water bath sonication. Lysates were then centrifuged for 10 min at 12,000 × g at 4°C. In experiments where cells were lysed in-well to assess total protein content, cells were lysed by adding protease inhibitor cocktail and Triton-X-100 1% (v/v) into the culture medium. In-well lysates were concentrated by mixing with an equal volume of trichloroacetic acid (Fisher, 10391351) and centrifuged for 10 min at 18,000 × g at 4°C. The supernatant was discarded, and the pellet resuspended in acetone (100%) before centrifugation for 10 min at 18,000 × g at 4°C. The supernatant was again removed and the pellet allowed to air dry, before resuspending in Laemmli buffer (2X). Samples were analysed for NRF2, pro-IL-1β, mature IL-1β, NLRP3, pro-caspase-1, caspase-1 p10, and gasdermin D. Equal amounts of protein from lysates or equal volumes of in-well lysates were loaded into the gel. Samples were run on SDS-polyacrylamide gels and transferred at 25 V onto nitrocellulose or PVDF membranes using a Trans-Blot^®^ Turbo Transfer™ System (Bio-Rad). The membranes were blocked in either 5% w/v milk or 2.5% BSA (Sigma, A3608) in PBS, 0.1% Tween 20 (PBST) for 1 h at room temperature. The membranes were then washed with PBST and incubated at 4°C overnight with goat anti-mouse IL-1β (250 ng ml^−1^; R&D Systems, AF-401-NA), mouse anti-mouse NLRP3 (1 μg ml^−1^; Adipogen, G-20B-0014-C100), rabbit anti-mouse caspase-1 (1.87 μg ml^−1^; Abcam, ab179515), rabbit anti-mouse gasdermin D (0.6 μg ml^−1^; Abcam, ab209845) or rabbit anti-mouse NRF2 (1.5 μg ml^−1^; CST, 12721) primary antibodies in 0.1% (IL-1β), 1% (NLRP3) or 2.5% (caspase-1, gasdermin D, NRF2) BSA in PBST. The membranes were washed and incubated with rabbit anti-goat IgG (500 ng ml^−1^, 5% milk in PBST; Dako, P044901-2), rabbit anti-mouse IgG (1.3 μg ml^−1^, 5% milk in PBST; Dako, P026002-2) or goat anti-rabbit IgG (250 ng ml^−1^, 2.5% BSA in PBST; Dako, (Dako, P044801-2) at room temperature for 1 h. Proteins were then visualised with Amersham ECL Western Blotting Detection Reagent (GE Healthcare, RPN2236) and G:BOX (Syngene) and Genesys software. β-Actin (Sigma, A3854) was used as a loading control. Densitometry was performed using FIJI (ImageJ). Uncropped western blots are provided in Supplementary Figures 7-11.

### ELISA

The levels of IL-1β, IL-6 and tumour necrosis factor (TNF) in the supernatant were analysed by enzyme-linked immunosorbent assay (ELISA; DuoSet, R&D systems) according to the manufacturer’s instructions.

### Cell death assays

Cell death was assessed by measuring lactate dehydrogenase (LDH) release into the supernatant using a CytoTox 96 Non-Radioactive Cytotoxicity Assay (Promega) according to the manufacturer’s instructions. Cell death in OHSCs was assessed by adding propidium iodide (25 μg ml^−1^; Sigma, P4864) to the culture medium for the final 30 min of the inflammasome activation protocol followed by widefield microscopy.

### Live imaging of ASC speck formation

ASC–citrine-expressing primary BMDMs were used to perform live imaging of ASC speck formation. For itaconate derivative pre-treatment assays, cells were seeded at 1 × 10^6^ cells ml^−1^ in 96-well plates and incubated for 1 h, and were then treated with vehicle (DMSO), DMI, 4OI or DMF (125 μM, 20 h). LPS (1 μg ml^−1^, 4 h) was then added to the wells to induce priming. The medium was replaced with optimem, and nigericin (10 μM) was added to activate the NLRP3 inflammasome. For assays where itaconate derivative treatments were added after LPS priming, cells were seeded overnight at 1 × 10^6^ cells ml^−1^ in 96-well plates. Cells were then first primed with LPS (1 μg ml^−1^, 4 h). The medium was replaced with optimum containing vehicle, DMI, 4OI or DMF (125 μM, 15 min) prior to addition of nigericin (10 μM). Image acquisition began immediately after nigericin treatment. Images were subsequently acquired every 10 min for a further 90 min using an IncuCyte ZOOM^®^ Live Cell Analysis system (Essen Bioscience) at 37 °C using a 20X/0.61 S Plan Fluor objective. Speck number was quantified using IncuCyte ZOOM^®^ software, and was assessed for each treatment at the final time point of 90 min.

### OHSC immunostaining

OHSCs were washed once with cold PBS and fixed in 4% paraformaldehyde (1 h) at 4°C. OHSCs were washed two more times in cold PBS and then incubated with rabbit anti-mouse ASC (202 ng ml^−1^; CST, 67824) primary antibody overnight at 4°C. OHSCs were washed and incubated with Alexa Fluor^™^ 488 donkey anti-rabbit IgG (2 μg ml^−1^; Invitrogen, A-21206) secondary antibody for 2 h at room temperature. All antibody incubations were performed using PBS, 0.3% Triton X-100. Wash steps were performed using PBST unless stated otherwise. OHSCs were washed and then incubated in DAPI (1 μg ml^−1^, 15 min; Sigma, D9542) at room temperature before final washing and mounting using ProLong™ gold antifade mountant (Thermo, P36934) prior to imaging using widefield microscopy.

### Snapshot widefield microscopy

Images were collected on a Zeiss Axioimager.M2 upright microscope using a 5X or 20X Plan Apochromat objective and captured using a Coolsnap HQ2 camera (Photometrics) through Micromanager software (v1.4.23). Specific band-pass filter sets for DAPI and FITC were using to prevent bleed-through from one channel to the next.

### Image processing analysis

Analysis was performed using FIJI (ImageJ) on images acquired from the same region of up to three separate OHSCs (from the same insert) per treatment, and these values were averaged for each biological repeat. ASC speck formation was quantified on 20X widefield microscopy images by subtracting background (50 pixel rolling ball radius), manually setting thresholds and analysing particles with the following parameters: size 1–10 μm^2^, circularity 0.9–1.0. To quantify PI uptake, images were acquired on a widefield microscope using a 5X objective, background was subtracted (5.0 pixel rolling ball radius) and thresholds for images were automatically determined using the default method. The total area of PI-positive signal was measured in the whole field of view, and was then normalised to the total area of DAPI signal.

## Data analysis

Data are presented as the mean ± standard error of the mean (SEM) together with individual data points where possible. Data were analysed using repeated-measures one-way or two-way analysis of variance (ANOVA), or mixed effects model, with Dunnett’s or Sidak’s post-hoc test using GraphPad Prism (v8). Transformations or corrections were applied as necessary to obtain equal variance between groups prior to analysis. Statistical significance was accepted at *P<0.05.

## 5 Acknowledgments

ASC–citrine mice were a kind gift from Douglas Golenbock (University of Massachusetts Medical School) and Te-Chen Tzeng (Bristol Myers-Squibb, Cambridge, USA). We thank Dr Kevin Stacey (University of Manchester) for the routine isolation of human PBMCs.

## 8 Data availability

The data that support the findings of this study are available from the corresponding author upon reasonable request.

## 9 Author Contributions

Conceptualisation: CH, EL, SMA and DB. Methodology: CH, EL and DB. Investigation: CH, EL and JPG. Writing - original draft preparation: CH. Writing - review and editing: CH, EL, SMA and DB. Visualisation: CH. Supervision: EL, SMA and DB. Funding acquisition: DB and SMA.

## References

Bambouskova, M., Gorvel, L., Lampropoulou, V., Sergushichev, A., Loginicheva, E., Johnson, K., Korenfeld, D., Mathyer, M. E., Kim, H., Huang, L. H., et al. (2018). Electrophilic properties of itaconate and derivatives regulate the IκBζ-ATF3 inflammatory axis. Nature 556, 501–504.

Boucher, D., Monteleone, M., Coll, R. C., Chen, K. W., Ross, C. M., Teo, J. L., Gomez, G. A., Holley, C. L., Bierschenk, D., Stacey, K. J., et al. (2018). Caspase-1 self-cleavage is an intrinsic mechanism to terminate inflammasome activity. J. Exp. Med. 215, 827–840.

Chausse, B., Lewen, A., Poschet, G. and Kann, O. (2020). Selective inhibition of mitochondrial respiratory complexes controls the transition of microglia into a neurotoxic phenotype in situ. Brain. Behav. Immun. S0889-1591, 30209–9.

Chen, J. and Chen, Z. J. (2018). PtdIns4P on dispersed trans-Golgi network mediates NLRP3 inflammasome activation. Nature 564, 71–76.

Chen, G. Y. and Nuñez, G. (2010). Sterile inflammation: Sensing and reacting to damage. Nat. Rev. Immunol. 10, 826–837.

Coll, R. C., Robertson, A. A., Chae, J. J., Higgins, S. C., Muñoz-Planillo, R., Inserra, M. C., Vetter, I., Dungan, L. S., Monks, B. G., Stutz, A., et al. (2015). A small molecule inhibitor of the NLRP3 inflammasome for the treatment of inflammatory diseases. Nat Med 21, 248–255.

Cunha, M. I., Su, M., Cantuti-Castelvetri, L., Müller, S. A., Schifferer, M., Djannatian, M., Alexopoulos, I., van der Meer, F., Winkler, A., van Ham, T. J., et al. (2020). Pro-inflammatory activation following demyelination is required for myelin clearance and oligodendrogenesis. J. Exp. Med. 217, e20191390.

Dostert, C., Pétrilli, V., Van Bruggen, R., Steele, C., Mossman, B. T. and Tschopp, J. (2008). Innate Immune Activation Through Nalp3 Inflammasome Sensing of Asbestos and Silica. Science. 320, 674–677.

ElAzzouny, M., Tom, C. T. M. B., Evans, C. R., Olson, L. L., Tanga, M. J., Gallagher, K. A., Martin, B. R. and Burant, C. F. (2017). Dimethyl itaconate is not metabolized into itaconate intracellularly. J. Biol. Chem. 292, 4766–4769.

Fox, R. J., Miller, D. H., Phillips, J. T., Hutchinson, M., Havrdova, E., Kita, M., Yang, M., Raghupathi, K., Novas, M., Sweetser, M. T., et al. (2012). Placebo-Controlled Phase 3 Study of Oral BG-12 or Glatiramer in Multiple Sclerosis. N. Engl. J. Med. 367, 1087–1097.

Freeman, L., Guo, H., David, C. N., Brickey, W. J., Jha, S. and Ting, J. P. Y. (2017). NLR members NLRC4 and NLRP3 mediate sterile inflammasome activation in microglia and astrocytes. J. Exp. Med. 214, 1351–1370.

Gaidt, M. M., Ebert, T. S., Chauhan, D., Schmidt, T., Schmid-Burgk, J. L., Rapino, F., Robertson, A. A. B., Cooper, M. A., Graf, T. and Hornung, V. (2016). Human Monocytes Engage an Alternative Inflammasome Pathway. Immunity 44, 833–846.

Garstkiewicz, M., Strittmatter, G. E., Grossi, S., Sand, J., Fenini, G., Werner, S., French, L. E. and Beer, H. D. (2017). Opposing effects of Nrf2 and Nrf2-activating compounds on the NLRP3 inflammasome independent of Nrf2-mediated gene expression. Eur. J. Immunol. 47, 806–817.

Gold, R., Linker, R. A. and Stangel, M. (2012a). Fumaric acid and its esters: An emerging treatment for multiple sclerosis with antioxidative mechanism of action. Clin. Immunol. 142, 44–48.

Gold, R., Kappos, L., Arnold, D. L., Bar-Or, A., Giovannoni, G., Selmaj, K., Tornatore, C., Sweetser, M. T., Yang, M., Sheikh, S. I., et al. (2012b). Placebo-Controlled Phase 3 Study of Oral BG-12 for Relapsing Multiple Sclerosis. N. Engl. J. Med. 367, 1098–1107.

Gris, D., Ye, Z., Iocca, H. A., Wen, H., Craven, R. R., Gris, P., Huang, M., Schneider, M., Miller, S. D. and Ting, J. P.-Y. (2010). NLRP3 plays a critical role in the development of experimental autoimmune encephalomyelitis by mediating Th1 and Th17 responses. J. Immunol. 185, 974–81.

Groß, C. J., Mishra, R., Schneider, K. S., Médard, G., Wettmarshausen, J., Dittlein, D. C., Shi, H., Gorka, O., Koenig, P.-A., Fromm, S., et al. (2016). K+ Efflux-Independent NLRP3 Inflammasome Activation by Small Molecules Targeting Mitochondria. Immunity 45, 761–773.

Hall, S. M. (1972). The Effect of Injections of Lysophosphatidyl Choline into White Matter of the Adult Mouse Spinal Cord. J. Cell Sci. 10,.

Halle, A., Hornung, V., Petzold, G. C., Stewart, C. R., Monks, B. G., Reinheckel, T., Fitzgerald, K. A., Latz, E., Moore, K. J. and Golenbock, D. T. (2008). The NALP3 inflammasome is involved in the innate immune response to amyloid-beta. Nat Immunol 9, 857–865.

He, W.-T., Wan, H., Hu, L., Chen, P., Wang, X., Huang, Z., Yang, Z.-H., Zhong, C.-Q. and Han, J. (2015). Gasdermin D is an executor of pyroptosis and required for interleukin-1β secretion. Cell Res. 25, 1285–98.

Hooftman, A., Angiari, S., Hester, S., Corcoran, S. E., Runtsch, M. C., Ling, C., Ruzek, M. C., Slivka, P. F., McGettrick, A. F., Banahan, K., et al. (2020). The Immunomodulatory Metabolite Itaconate Modifies NLRP3 and Inhibits Inflammasome Activation. Cell Metab. 32, 468–478.e7.

Hornung, V., Bauernfeind, F., Halle, A., Samstad, E. O., Kono, H., Rock, K. L., Fitzgerald, K. A. and Latz, E. (2008). Silica crystals and aluminum salts activate the NALP3 inflammasome through phagosomal destabilization. Nat. Immunol. 9, 847–856.

Hoyle, C., Redondo‐Castro, E., Cook, J., Tzeng, T., Allan, S. M., Brough, D. and Lemarchand, E. (2020). Hallmarks of NLRP3 inflammasome activation are observed in organotypic hippocampal slice culture. Immunology imm.13221.

Humphries, F., Shmuel-Galia, L., Ketelut-Carneiro, N., Li, S., Wang, B., Nemmara, V. V., Wilson, R., Jiang, Z., Khalighinejad, F., Muneeruddin, K., et al. (2020). Succination inactivates gasdermin D and blocks pyroptosis. Science (80-.). 369, 1633–1637.

Jha, S., Srivastava, S. Y., Brickey, W. J., Iocca, H., Toews, A., Morrison, J. P., Chen, V. S., Gris, D., Matsushima, G. K. and Ting, J. P. Y. (2010). The inflammasome sensor, NLRP3, regulates CNS inflammation and demyelination via caspase-1 and interleukin-18. J. Neurosci. 30, 15811–15820.

Kayagaki, N., Warming, S., Lamkanfi, M., Walle, L. Vande, Louie, S., Dong, J., Newton, K., Qu, Y., Liu, J., Heldens, S., et al. (2011). Non-canonical inflammasome activation targets caspase-11. Nature 479, 117–121.

Kobayashi, E. H., Suzuki, T., Funayama, R., Nagashima, T., Hayashi, M., Sekine, H., Tanaka, N., Moriguchi, T., Motohashi, H., Nakayama, K., et al. (2016). Nrf2 suppresses macrophage inflammatory response by blocking proinflammatory cytokine transcription. Nat. Commun. 7, 11624.

Kornberg, M. D., Bhargava, P., Kim, P. M., Putluri, V., Snowman, A. M., Putluri, N., Calabresi, P. A. and Snyder, S. H. (2018). Dimethyl fumarate targets GAPDH and aerobic glycolysis to modulate immunity. Science (80-.). 360, 449–453.

Kuo, P. C., Weng, W. T., Scofield, B. A., Paraiso, H. C., Brown, D. A., Wang, P. Y., Yu, I. C. and Yen, J. H. (2020a). Dimethyl itaconate, an itaconate derivative, exhibits immunomodulatory effects on neuroinflammation in experimental autoimmune encephalomyelitis. J. Neuroinflammation 17,.

Kuo, P.-C., Weng, W.-T., Scofield, B. A., Furnas, D., Paraiso, H. C., Yu, I.-C. and Yen, J.-H. (2020b). Induction of IRG1 following ischemic stroke promotes HO-1 and BDNF expression to alleviate neuroinflammation and restrain ischemic brain injury. J. Neuroinflammation.

Lassmann, H. and Bradl, M. (2017). Multiple sclerosis: experimental models and reality. Acta Neuropathol. 133, 223–244.

Law, S. H., Chan, M. L., Marathe, G. K., Parveen, F., Chen, C. H. and Ke, L. Y. (2019). An updated review of lysophosphatidylcholine metabolism in human diseases. Int. J. Mol. Sci. 20, 1149.

Lemarchand, E., Barrington, J., Chenery, A., Haley, M., Coutts, G., Allen, J. E., Allan, S. M. and Brough, D. (2019). Extent of Ischemic Brain Injury After Thrombotic Stroke Is Independent of the NLRP3 (NACHT, LRR and PYD Domains-Containing Protein 3) Inflammasome. Stroke 50, 1232–1239.

Li, S., Wu, Y., Yang, D., Wu, C., Ma, C., Liu, X., Moynagh, P. N., Wang, B., Hu, G. and Yang, S. (2019). Gasdermin D in peripheral myeloid cells drives neuroinflammation in experimental autoimmune encephalomyelitis. J. Exp. Med. 216, 2562–2581.

Liao, S. T., Han, C., Xu, D. Q., Fu, X. W., Wang, J. S. and Kong, L. Y. (2019). 4-Octyl itaconate inhibits aerobic glycolysis by targeting GAPDH to exert anti-inflammatory effects. Nat. Commun. 10, 5091.

Linker, R. A., Lee, D. H., Ryan, S., Van Dam, A. M., Conrad, R., Bista, P., Zeng, W., Hronowsky, X., Buko, A., Chollate, S., et al. (2011). Fumaric acid esters exert neuroprotective effects in neuroinflammation via activation of the Nrf2 antioxidant pathway. Brain 134, 678–692.

Litjens, N. H. R., Burggraaf, J., Van Strijen, E., Van Gulpen, C., Mattie, H., Schoemaker, R. C., Van Dissel, J. T., Thio, H. B. and Nibbering, P. H. (2004). Pharmacokinetics of oral fumarates in healthy subjects. Br. J. Clin. Pharmacol. 58, 429–432.

Liu, X., Zhou, W., Zhang, X., Lu, P., Du, Q., Tao, L., Ding, Y., Wang, Y. and Hu, R. (2016). Dimethyl fumarate ameliorates dextran sulfate sodium-induced murine experimental colitis by activating Nrf2 and suppressing NLRP3 inflammasome activation. Biochem. Pharmacol. 112, 37–49.

Liu, L., Locascio, L. M. and Doré, S. (2019). Critical Role of Nrf2 in Experimental Ischemic Stroke. Front. Pharmacol. 10, 153.

McKenzie, B. A., Mamik, M. K., Saito, L. B., Boghozian, R., Monaco, M. C., Major, E. O., Lu, J. Q., Branton, W. G. and Power, C. (2018). Caspase-1 inhibition prevents glial inflammasome activation and pyroptosis in models of multiple sclerosis. Proc. Natl. Acad. Sci. U. S. A. 115, E6065–E6074.

Miglio, G., Veglia, E. and Fantozzi, R. (2015). Fumaric acid esters prevent the NLRP3 inflammasome-mediated and ATP-triggered pyroptosis of differentiated THP-1 cells. Int. Immunopharmacol. 28, 215–219.

Mills, E. and O’Neill, L. A. J. (2014). Succinate: A metabolic signal in inflammation. Trends Cell Biol. 24, 313–320.

Mills, E. L., Ryan, D. G., Prag, H. a., Dikovskaya, D., Menon, D., Zaslona, Z., Jedrychowski, M. P., Costa, A. S. H., Higgins, M., Hams, E., et al. (2018). Itaconate is an anti-inflammatory metabolite that activates Nrf2 via alkylation of KEAP1. Nature 556, 113–117.

Muñoz-Planillo, R., Kuffa, P., Martínez-Colón, G., Smith, B., Rajendiran, T. and Núñez, G. (2013). K+ Efflux Is the Common Trigger of NLRP3 Inflammasome Activation by Bacterial Toxins and Particulate Matter. Immunity 38, 1142–1153.

Nair, S., Huynh, J. P., Lampropoulou, V., Loginicheva, E., Esaulova, E., Gounder, A. P., Boon, A. C. M., Schwarzkopf, E. A., Bradstreet, T. R., Edelson, B. T., et al. (2018). Irg1 expression in myeloid cells prevents immunopathology during M. tuberculosis infection. J. Exp. Med. 215, 1035–1045.

O’Neill, L. A. J. and Artyomov, M. N. (2019). Itaconate: the poster child of metabolic reprogramming in macrophage function. Nat. Rev. Immunol. 19, 273–281.

Olagnier, D., Farahani, E., Thyrsted, J., Blay-Cadanet, J., Herengt, A., Idorn, M., Hait, A., Hernaez, B., Knudsen, A., Iversen, M. B., et al. (2020). SARS-CoV2-mediated suppression of NRF2-signaling reveals potent antiviral and anti-inflammatory activity of 4-octyl-itaconate and dimethyl fumarate. Nat. Commun. 11, 4938.

Perregaux, D. and Gabel, C. a. (1994). Interleukin-1β maturation and release in response to ATP and nigericin. Evidence that potassium depletion mediated by these agents is a necessary and common feature of their activity. J. Biol. Chem. 269, 15195–15203.

Pinteaux, E., Parker, L. C., Rothwell, N. J. and Luheshi, G. N. (2002). Expression of interleukin-1 receptors and their role in interleukin-1 actions in murine microglial cells. J. Neurochem. 83, 754–763.

Plemel, J. R., Michaels, N. J., Weishaupt, N., Caprariello, A. V., Keough, M. B., Rogers, J. A., Yukseloglu, A., Lim, J., Patel, V. V., Rawji, K. S., et al. (2018). Mechanisms of lysophosphatidylcholine-induced demyelination: A primary lipid disrupting myelinopathy. Glia 66, 327–347.

Qin, W., Zhang, Y., Tang, H., Liu, D., Chen, Y., Liu, Y. and Wang, C. (2020). Chemoproteomic Profiling of Itaconation by Bioorthogonal Probes in Inflammatory Macrophages. J. Am. Chem. Soc. 142, 10894–10898.

Rock, K. L., Latz, E., Ontiveros, F. and Kono, H. (2010). The sterile inflammatory response. Annu. Rev. Immunol. 28, 321–42.

Schaum, N., Karkanias, J., Neff, N. F., May, A. P., Quake, S. R., Wyss-Coray, T., Darmanis, S., Batson, J., Botvinnik, O., Chen, M. B., et al. (2018). Single-cell transcriptomics of 20 mouse organs creates a Tabula Muris. Nature 562, 367–372.

Schroder, K. and Tschopp, J. (2010). The Inflammasomes. Cell 140, 821–832.

Schulze-Topphoff, U., Varrin-Doyer, M., Pekarek, K., Spencer, C. M., Shetty, A., Sagan, S. A., Cree, B. A. C., Sobel, R. A., Wipke, B. T., Steinman, L., et al. (2016). Dimethyl fumarate treatment induces adaptive and innate immune modulation independent of Nrf2. Proc. Natl. Acad. Sci. U. S. A. 113, 4777–4782.

Seoane, P. I., Lee, B., Hoyle, C., Yu, S., Lopez-Castejon, G., Lowe, M. and Brough, D. (2020). The NLRP3-inflammasome as a sensor of organelle dysfunction. J. Cell Biol. 219, e202006194.

Sheppard, O., Coleman, M. P. and Durrant, C. S. (2019). Lipopolysaccharide-induced neuroinflammation induces presynaptic disruption through a direct action on brain tissue involving microglia-derived interleukin 1 beta. J. Neuroinflammation 16, 1–13.

Smith, D. (2017). Fumaric acid esters for psoriasis: a systematic review. Ir. J. Med. Sci. 186, 161–177.

Stoppini, L., Buchs, P. and Muller, D. (1991). A simple method for organotypic cultures of nervous tissue. J. Neurosci. Methods 37, 173–182.

Swain, A., Bambouskova, M., Kim, H., Andhey, P. S., Duncan, D., Auclair, K., Chubukov, V., Simons, D. M., Roddy, T. P., Stewart, K. M., et al. (2020). Comparative evaluation of itaconate and its derivatives reveals divergent inflammasome and type I interferon regulation in macrophages. Nat. Metab. 1–9.

Tannahill, G., Curtis, A., Adamik, J., Palsson-McDermott, E., McGettrick, A., Goel, G., Frezza, C., Bernard, N., Kelly, B., Foley, N., et al. (2013). Succinate is an inflammatory signal that induces IL-1β via HIF-1α. Nature 496, 238–242.

Trotter, A., Anstadt, E., Clark, R. B., Nichols, F., Dwivedi, A., Aung, K. and Cervantes, J. L. (2019). The role of phospholipase A2 in multiple Sclerosis: A systematic review and meta-analysis. Mult. Scler. Relat. Disord. 27, 206–213.

Tzeng, T. C., Schattgen, S., Monks, B., Wang, D., Cerny, A., Latz, E., Fitzgerald, K. and Golenbock, D. T. (2016). A Fluorescent Reporter Mouse for Inflammasome Assembly Demonstrates an Important Role for Cell-Bound and Free ASC Specks during In Vivo Infection. Cell Rep. 16, 571–582.

Wilms, H., Sievers, J., Rickert, U., Rostami-Yazdi, M., Mrowietz, U. and Lucius, R. (2010). Dimethylfumarate inhibits microglial and astrocytic inflammation by suppressing the synthesis of nitric oxide, IL-1β, TNF-α and IL-6 in an in-vitro model of brain inflammation. J. Neuroinflammation 7, 30.

Zhang, Y., Chen, K., Sloan, S. A., Bennett, M. L., Scholze, A. R., O’Keeffe, S., Phatnani, H. P., Guarnieri, P., Caneda, C., Ruderisch, N., et al. (2014). An RNA-Sequencing Transcriptome and Splicing Database of Glia, Neurons, and Vascular Cells of the Cerebral Cortex. J. Neurosci. 34, 11929–11947.

